# Bacterial surface display enables lysis-independent joint host–pathogen single-cell profiling

**DOI:** 10.64898/2026.07.21.739887

**Authors:** Cal Gunnarsson, Jeff C. Hsiao, Jacob B. Hochfelder, Sydney L. Solomon, Bianca A. Lepe, Sharon Zhu, Kelly Xu, Hattie A. Chung, Bryan D. Bryson

## Abstract

Heterogeneity in host–pathogen interactions arises from variation in both host cell state and pathogen state, yet most single-cell methods capture these features separately. Joint profiling is limited by fundamental technical mismatches between host- and pathogen-derived material, particularly differences in cell wall structure, lysis requirements, and molecular abundance. Here we introduce a lysis-independent strategy that enables unified measurement of intracellular bacterial presence and state alongside host single-cell profiles. We repurpose bacterial surface display to encode promoter activity and bacterial identity as antibody-detectable signals, rendering bacterial features compatible with existing antibody-based single-cell assays. This approach is modular across promoters, epitope tags, display scaffolds, and bacterial species, including *Escherichia coli* and *Mycobacterium tuberculosis*. Surface-displayed reporters are detectable during intracellular infection and can be read out by flow cytometry and droplet-based single-cell RNA sequencing without pathogen-specific lysis optimization or pre-sorting on pathogen signal. Applying this method to infected macrophages, we link heterogeneous bacterial uptake to heterogeneous expression of phagocytosis-associated host programs. This strategy enables scalable, joint host–pathogen single-cell measurements and expands the range of pathogens and states accessible to high-throughput single-cell analysis.

## Introduction

Hosts, pathogens, and their interactions are heterogeneous at the single-cell level, driving heterogeneous infection and treatment outcomes. Host cells of the same cell type and source differ in phagocytic,^1^ antimicrobial,^2–4^ and cytokine production capacities.^5–7^ Intracellular bacteria form stressed,^8^ stress-resistant, and antibiotic-resistant subpopulations,^3,9–12^ and heterogeneous bacterial cell surfaces lead to heterogeneous host sensing.^13^ Besides potentially existing pre-infection, heterogeneity is often an emergent property driven by host-pathogen feedback. Measuring heterogeneity, identifying its sources, and defining strategies for therapeutic manipulation is aided by a method’s coverage of both host and pathogen.

Single-cell methods, particularly single-cell RNA sequencing (scRNAseq),^6,14–22^ have mapped host and pathogen heterogeneity separately. However, fewer methods link information about pathogens and the single cells they infect. Existing highly multiplexable methods rely on detecting native^23,24^ or engineered^25,26^ pathogen cytosolic species, adding technical requirements related to cytosolic access and single-cell scales. Highly multiplexable methods require controlled compartmentalization of single cells, either creating single-cell–containing compartments where material is extracted and uniquely labeled per-cell or assaying gently permeabilized intact cells. These requirements are sensitive to variations in cell physiology present across the tree of life. Unicellular pathogens have thick cell walls with diverse compositions, less cellular material, and may lack enrichable molecular modifications (e.g. mRNA polyadenylation, for prokaryotes). Accordingly, methods developed for mammalian cells often yield little material when directly applied to pathogens.^14,19^ Extensive species- and method-specific optimization is required to access pathogen material, as extraction methods do not necessarily translate within a microbial kingdom, and harsher extraction is generally at odds with maintenance of single-cell compartments, especially for high-throughput droplet-based and split-pool methods.^14,19^ For joint host-pathogen profiling, dissimilarity between host and pathogen cell membranes introduces challenges for single-cell compartmentalization, and difficulty extracting pathogen-derived material is exacerbated by abundant host-derived material.

Removing the requirement for lysis would obviate specialized optimization, but lysis-free methods typically profile features in a more limited set of host-focused assays. Classically, transcriptional reporters for intracellular pathogens use bright fluorophores to allow lysis-free measurement of single-cell state, but are limited by spectral overlap and limited to optical assays or experiments on sorted subpopulations. Bridging the gap between fluorescent transcriptional reporting and specialized, highly multiplexable methods, antibody-based epitope detection allows high-dimensional measurement of native proteins, epitopes, or epitope combinations^27,28^ and is directly compatible with multiple detection modalities.^29–32^ As epitopes are detectable across modalities, antibody-based strategies both simplify workflows by eliminating cell sorting and further resolve pathogen features that would otherwise be binned during sorting.

We describe surface display of linear epitopes as a modular strategy for reporting on bacterial state and presence, allowing antibody-based detection in existing multiplexed assays of host state without optimizing bacterial lysis, enrichment, or requiring cell sorting. Activity of a promoter of interest is linked to surface display of an epitope-tagged protein, rendering promoter activity as an antibody-detectable feature. We demonstrate this technique with axenic *Escherichia coli* (E.coli) and *Mycobacterium tuberculosis* (Mtb), define engineering principles for surface display, and characterize parameters governing display during intracellular infection. Using this technique, we detect bacterial-derived signal in infected cells by flow cytometry and by droplet-based scRNAseq. By linking bacterial-derived signal to host state, we show that heterogeneous uptake of *E.coli* is linked to heterogeneous expression of phagocytosis-regulatory genes.

## Results

### Surface display converts identity and promoter activity into antibody-detectable signal

Since yeast^33^ and bacterial surface display^34^ use inducible promoters to display a cargo of interest, we reasoned surface display could be adapted as a reporter for a given promoter (Figure 1A). In this paradigm, surface staining of an epitope-tagged display scaffold is a proxy for promoter activity, and specific epitope tags encode specific promoters. To establish this approach, we tagged the engineered surface display scaffold eCPX^35^ with a C-terminal hemagglutinin (HA) tag, then drove its expression in *E.coli* using an anhydrotetracycline (aTc)-responsive promoter, pTet (Figure 1D). This approach generalized to other promoters, including the native arabinose-inducible pBAD promoter and a constitutive promoter from the Anderson promoter library 105 (Figure 1D), other epitope tags (Figure 1B), and other scaffolds (Supplementary Figure 1A). E.coli strains expressing different epitope tags on their surface could only be stained with the appropriate cognate antibody (Figure 1B), and multiple epitope-tagged scaffolds could be simultaneously expressed and detected in single *E.coli* (Figure 1C). We additionally tested the use of this approach for reporting on the activity of bacterial promoters related to acid stress. We observe tag signal induction at pH 4.7 in E.coli expressing eCPX-His under the control acid-inducible pASR promoter (Figure 1E). This approach also generalizes to other bacterial species. We engineered the *Mycobacterium tuberculosis* (Mtb) protein PPE51 with a Myc tag under the control of the acid-stress promoter Rv1405c, and observed Myc signal in Mtb cultured at pH 5.7 (Figure 1F). Taken together, epitope tags can be used to encode multiple, orthogonal signals for multiple bacterial species.

**Figure 1.**
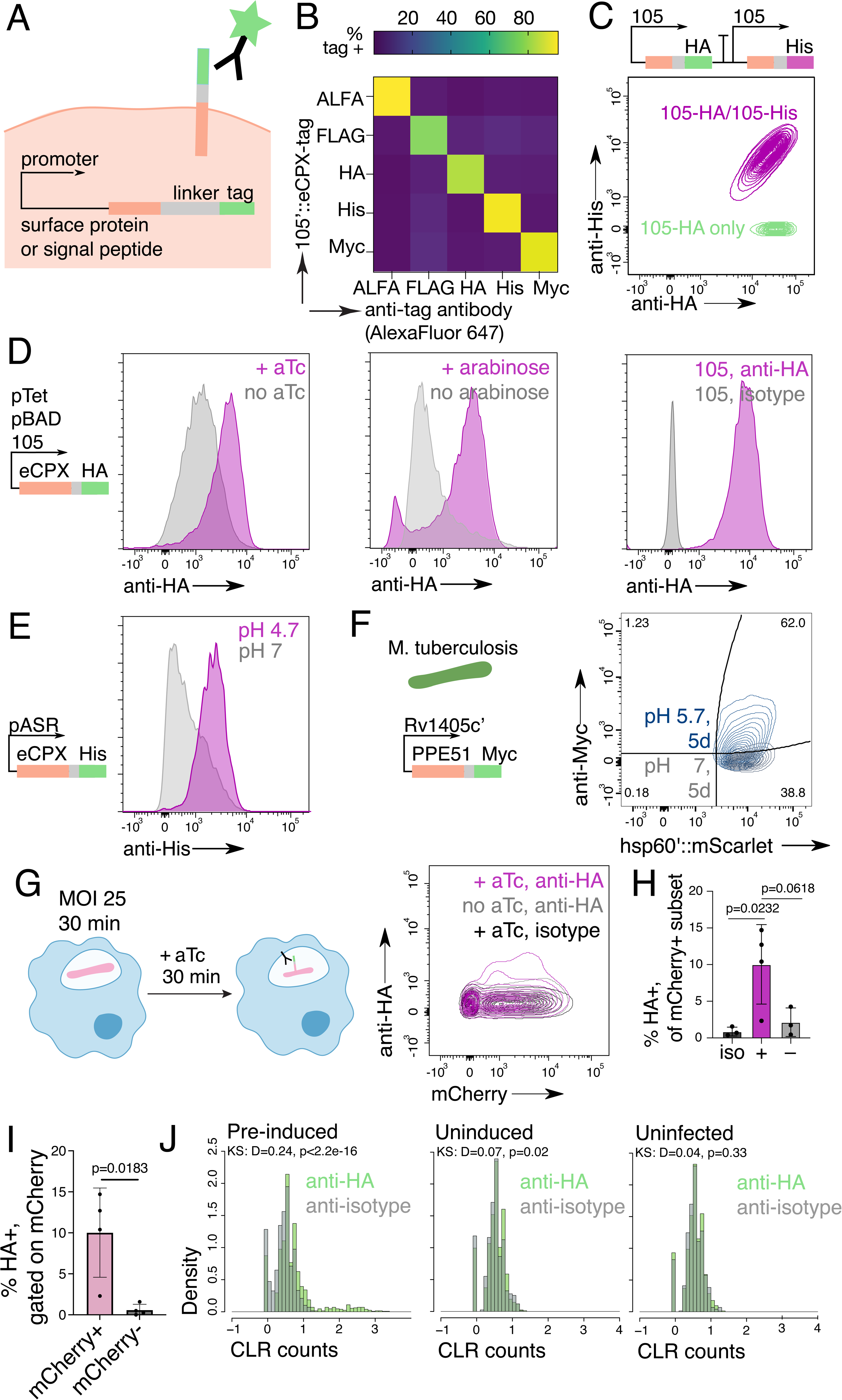
Surface display of a tagged surface protein links promoter activity to antibody-detectable signal during infection. **A)** Schematic of surface display. **B)** All-against-all staining of five strains constitutively expressing eCPX, each with different C-terminal tags. Representative of n=2 staining experiments performed on 2 different days. **C)** Display and staining of multiple epitope tags in single bacteria. Representative of n=2 staining experiments performed on 2 different days. **D)** Anhydrotetracycline-inducible (aTc), arabinose-inducible, and constitutive surface display using pTet, pBAD, and 105 promoters and C-terminally tagged eCPX. Representative of n=3 induction and staining experiments performed on 3 different days. **E)** Acid-inducible surface display using the native E. coli promoter pASR and His-tagged eCPX during culture in low pH or neutral LB for 1 hour. **F)** Surface staining of *Mycobacterium tuberculosis* surface protein PPE51. Acid inducible surface display using Rv1405c’ during culture in low pH or neutral minimal medium over 5 days. Representative of n=3 inductions performed on 2 different days. **G)** Intracellular HA staining and flow cytometry of infected iBMMs induced post-infection (+), relative to uninduced (-) and isotype-stained, aTc-induced conditions (iso). **H - I)** Statistical comparison of HA positivity in induced, uninduced, and isotype-stained directly infected (mCherry+) cells (**H**) or in directly infected (mCherry+) and bystander cells (mCherry-) in induced iBMMs (**I**); HA signal gated on isotype. **J)** Histogram of centered log ratio-normalized (CLR) HA- or isotype-derived counts in iBMMs infected with fixed, pre-induced (**H**) or fixed, uninduced E.coli (**I**) or uninfected iBMMs (**J**). Statistical analysis involving percentages were calculated based on the arcsin(sqrt(p)) transformation. Multiple comparisons were performed by one-way ANOVA with a post-hoc Tukey’s test for **H**. Statistical comparison of distributions in **J** were performed by two-sided Kolmogorov-Smirnov test. Error bars shown represent standard deviations.

Compared to transcriptional reporting by fluorescence, reporting by surface display requires export in addition to translation of a protein. To test whether the requirement for export delays signal detection, we induced a pTet::eCPX-HA display strain with aTc, then either fixed cells and stained for surface HA or lysed cells and blotted cell lysates for total HA at intervals along a 60-minute time course (Supplementary Figure 1B). Within the first 10 minutes of aTc addition, eCPX is expressed and becomes antibody-accessible (Supplementary Figure 1C). This timescale is similar to that of fluorescent protein maturation^36^ and suggests that surface display is not a rate-limiting step for reporting during mid-log growth. Similar trends were reflected in total protein lysates (Supplementary Figure 1D), where total HA was lower at earlier timepoints and correlated with surface HA signal (Supplementary Figure 1E). Other promoters (pBAD) and tags (His) exhibited similar surface display kinetics.

### Surface display enables detection of intracellular cargo across multiple antibody-based formats

We next tested whether surface display could be applied to report on bacterial promoter activity during intracellular infection. As proof-of-principle for intracellular bacterial reporting by surface display, we used the promoter pTet, as its small-molecule–responsive nature allowed us to define inactive and active conditions for the promoter. We transformed pTet::eCPX-HA into a constitutively fluorescent (pON::mCherry) E.coli strain,^37^ allowing us to orthogonally identify infected cells. We infected immortalized murine bone marrow macrophages (iBMMs) with live pTet::eCPX-HA / pON::mCherry E.coli, triple-washed iBMMs to remove extracellular bacteria, induced intracellular bacteria with aTc post-infection, and performed intracellular staining to detect bacterial-derived HA (Supplementary Figure 2A). In infected cells induced with aTc, HA signal was above isotype, demonstrating that signals derived from surface display strains are inducible and detectable during infection (Figure 1G-I). This signal was significantly higher than HA signal in aTc-treated bystander cells (Figure 1I) and non-aTc–treated, pTet::eCPX-HA–infected samples (Figure 1G), together showing that increased intracellular HA staining is a reporter of promoter induction, rather than an artefact of aTc addition or baseline leaky expression from pTet.

At high MOI, cells are more likely to be multiply infected, and detection of intracellular bacterial-derived epitopes could be aided by higher bacterial burden. To test the feasibility of detecting bacterial-derived epitopes at low MOI, we infected macrophages with pre-induced, fixed pTet::eCPX-HA/pON::mCherry E.coli at MOI 1, 5, 10, 25, and 50. Although percent HA positivity was slightly lower at MOI 1 compared to MOI 25, we did not observe a strong relationship between MOI and percent HA positivity, and percent HA positivity in induced samples was significantly higher than isotype stained and uninduced samples even at MOI 1 (Supplementary Figure 2B-C). Together, this suggests that reporting by surface display is applicable to low MOI infections and detecting single intracellular bacteria.

Through site-specific conjugation with fluorescent, oligonucleotide, metal, or enzyme moieties, antibody-based detection is itself modular, so we reasoned that reporting by surface display would be compatible with established multimodal, single cell RNA sequencing workflows. We infected iBMMs for 30 minutes with preinduced, fixed (eCPX-HA+) bacteria or uninduced, fixed (eCPX-HA-) bacteria, or performed a mock infection of uninfected iBMMs. After phagocytosis, we detached, fixed, permeabilized, and stained cells with DNA-conjugated HA and isotype antibodies, then prepared scRNAseq samples per standard droplet-based protocols. After quality control, we recovered 1,104, 653, and 603 cells in the induced, uninduced, and uninfected conditions, respectively. As expected for specific detection of bacterial-derived HA, sequencing-derived HA counts were above isotype counts in samples infected with pre-induced bacteria (Figure 2J), while distributions of HA counts closely matched the distribution of isotype-derived counts for uninduced and uninfected samples (Figure 2J). Together, we demonstrate that intracellular bacteria can be detected in host-focused single cell RNAseq using bacterial surface-displayed epitopes.

**Figure 2.**
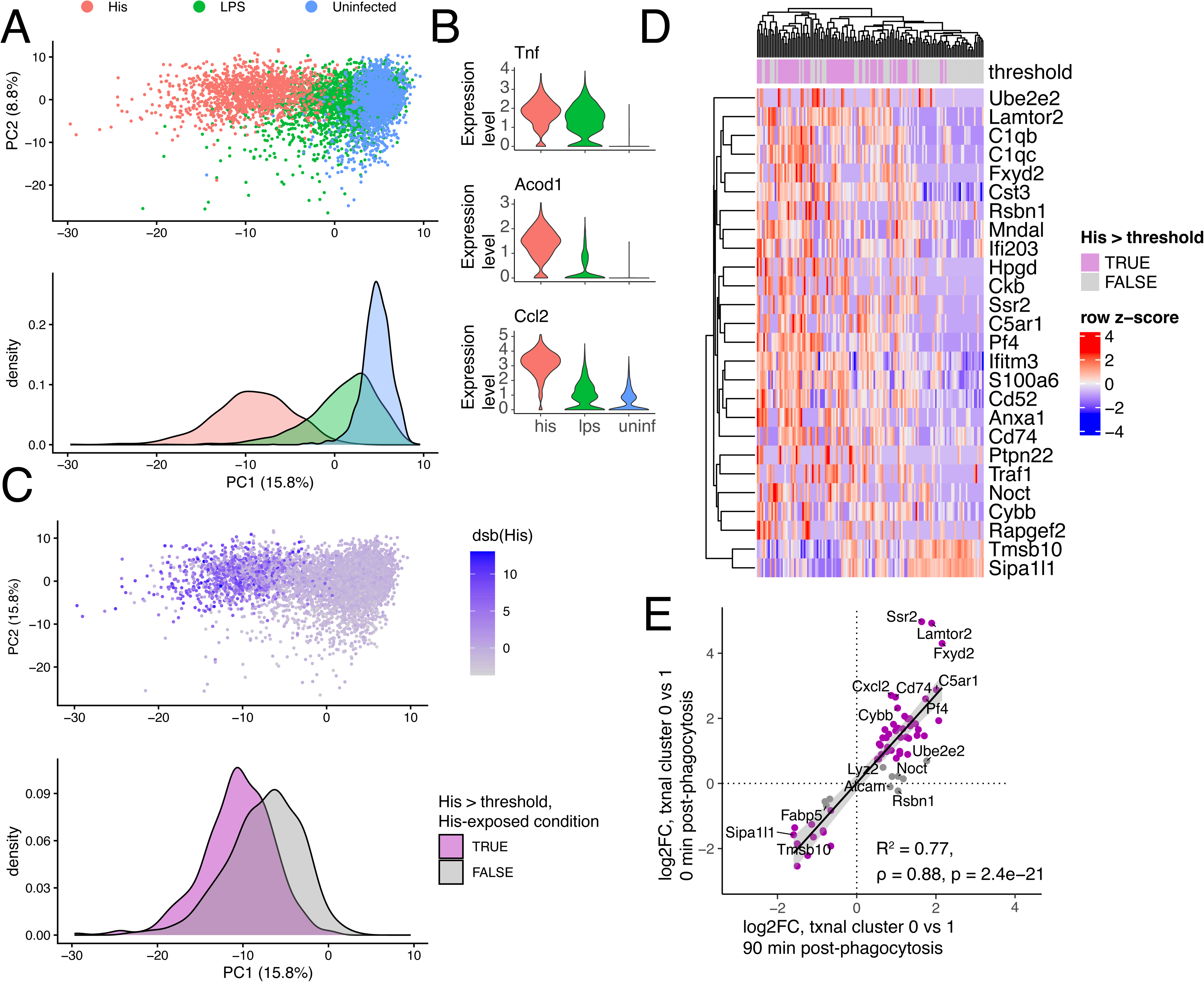
Joint measurement of bacterial presence and host state reveals a conserved signature of phagocytosis. **A)** Principal component projection of iBMMs infected with 108::eCPX-His *E.coli* at MOI 25 (His), treated with 100 ng/mL LPS (LPS), or uninfected at 90 minutes after a 30 minute phagocytosis period for infected samples and 120 minutes post-exposure for LPS-treated samples. Kernel density estimates for each condition are plotted along PC1. **B)** Scaled expression levels of Tnf, Acod1, and Ccl2 across conditions. **C)** Dsb-normalized His levels across conditions. **D)** Genes significantly associated with His levels within the His-infected condition are plotted for cells in the 5th and 95th quantiles of dsb(His expression). Genes are selected using stringent cutoffs (Methods). **E)** Comparison of transcriptional markers in 108::eCPX-His–infected samples at 90 minutes post-phagocytosis and Tet::eCPX-HA–infected samples at 0 minutes post-phagocytosis. Fold change between clusters is plotted for genes significantly associated with His levels using discovery cutoffs. Labels are shown for genes that meet stringent cutoffs. Correlations in **E** are calculated using linear regression and Pearson’s correlation test.

### Retention of bacteria-derived signal depends on epitope tag, phagosome state, and bacterial viability

Despite homogeneous display in the inoculum, cells infected with pre-fixed, HA-displaying bacteria did not homogeneously retain HA signal. Since HA contains a protease-cleavable motif,^38^ we tested whether loss of bacteria-derived signal was HA-specific by staining iBMMs infected with strains constitutively expressing mCherry and His- or HA-tagged eCPX (108::eCPX-HA/His / pON::mCherry) at 0, 30, 60, and 90 minutes post-phagocytosis. His positivity was retained up to 90 minutes post-phagocytosis (Supplementary Figure 3A-B), while percent HA positivity declined over time and was lower than percent His positivity at all matched timepoints.

Reporter signal stability influences time resolution and biological interpretation of positive and negative signals, so we next identified factors influencing HA loss. Because non-pathogenic E.coli are targeted to degradative compartments within the first hour of infection,^39^ we hypothesized that HA loss could be associated with loss of bacterial integrity. As a proxy for permeable, non-intact bacteria, we measured accessibility of mCherry in the bacterial cytosol using an anti-mCherry antibody (Supplementary Figure 3C-D). Antibody accessibility increased over time, and a majority of mCherry+ iBMMs stained positively for mCherry by 90 minutes post-phagocytosis. This timescale is consistent with orthogonal measurements of integrity loss in intracellular, non-pathogenic E.coli^40^ and suggests that His-tagged eCPX is retained even as bacterial cytosolic species become antibody-accessible. Consistent with a decrease in bacterial integrity and viability over time, inducing HA display later in infection led to proportionally fewer HA-positive bacteria at endpoint compared to inducing HA display early in infection (Supplementary Figure 4).

We next tested whether host degradative capacity influenced loss of HA, using bafilomycin A1 to inhibit phagosome maturation and acidification. Bafilomycin pretreatment almost entirely rescued aTc-responsiveness in live Tet::eCPX-HA E.coli (Supplementary Figure 3E-F). This suggests that loss of HA signal depends on phagosome state and bafilomycin-disrupted host processes. Taken together, we show that induction and retention of bacterial-derived signal depends on choice of epitope tag, phagosome state, and bacterial viability.

### Joint measurement of bacterial presence and host state recovers phagocytosis-regulatory host factors

Since His-expressing bacteria could be reliably detected intracellularly, we applied joint bacterial epitope and single cell RNA sequencing to identify host features associated with uptake of non-pathogenic *E.coli*. At 90 minutes post-infection, we detached, fixed, permeabilized, and stained iBMMs infected with 108::eCPX-His / pON::mCherry, as well as lipopolysaccharide (LPS)-treated and uninfected samples. After preparation and sequencing of gene expression and antibody-derived libraries, we recovered 1392, 2076, and 2349 high-quality cells, respectively. Although loss of complexity has been reported for libraries generated from fixed samples,^32^ we could nonetheless recover transcriptional differences between E.coli-infected, LPS-treated, and mock-infected samples in dimensionally reduced space (Figure 2A) and by differential gene expression (Supplementary Table 1; Figure 2B). E.coli-infected, LPS-treated, and mock-infected samples formed a continuum along a single principal component of transcriptional variation representing interferon-stimulated genes (ISG), such as Tnf, Acod1, and Ccl2 (Figure 2A-B; Supplementary Table 1). Stronger interferon responses in E.coli-exposed, compared to LPS-treated, samples may be due to differences in effective concentration of LPS delivered, activation of endosomal signaling, or response to pathogen-derived components in addition to LPS.

We next asked whether bacterial uptake, as indicated by His-derived antibody counts, was associated with a transcriptional signature. While His-positive cells did not form a discrete cluster in principal-component–reduced space, they were biased toward negative PC1 coordinates, suggesting an association between gene expression and bacterial uptake (Figure 2C). To nominate uptake-associated genes, we fit a generalized linear model (GLM) with raw counts of a given gene as output and normalized His counts, as well as technical and biological covariates, as input (Methods). Significant genes included Sipa1l1,^41^ Lamtor2,^42–44^ Ifitm3,^45^ Cd52,^43,46^ Ckb,^47^ and Anxa1^48–50^, which have previously described associations with or regulatory roles in phagocytosis (Figure 2D; Supplementary Table 1).

Using genes associated with bacterial-derived His and bacterial uptake, we examined whether macrophages exhibited phagocytosis-associated transcriptional heterogeneity at earlier timepoints. Immediately post-phagocytosis, cells separated into two transcriptional clusters in dimensionally reduced space, whether infected with eCPX::HA-displaying bacteria, infected with uninduced bacteria, or mock infected (Supplementary Figure 5A-C). Similar to His-positive cells at later timepoints, HA-positive cells exhibited skewed cluster membership, mapping to cluster 0 at greater frequency, and markers distinguishing the two clusters included genes previously identified as His-associated (Figure 2E; Supplementary Figure 5D). Marker expression was highly correlated between infected and uninfected samples, whether considering His-associated genes or all significant cluster markers (Supplementary Figure 5E-F), demonstrating that host heterogeneity is not attributable to interaction with surface-displayed eCPX and exists at early timepoints, including pre-infection. Reanalysis of two independently generated scRNAseq datasets supported heterogeneous expression of His-associated genes in unstimulated bone marrow-derived mouse macrophages, as well as stimulated mouse macrophages (Supplementary Figure 6A-B). Taken together, we jointly profile host state with bacterial presence, recovering host factors associated with phagocytosis that are heterogeneously expressed in multiple macrophage contexts.

### Surface display in non-model organisms is modular and tuned by export machinery availability

We next extended axenic surface display to the difficult-to-lyse bacterium *Mycobacterium tuberculosis* (Mtb). In Mtb, there are no known consensus signal sequences controlling outer membrane localization, and few known outer membrane proteins outside of ESX-5-secreted PPE proteins.^51^ Accordingly, we prototyped surface display using PPE protein scaffolds.

Since surface staining of C-terminally His-tagged PPE51 has been previously demonstrated,^52^ we tested whether PPE51 surface display was also retained upon C-terminal tagging with other epitope tags. PPE51 could be tagged and detected with multiple epitope tag-antibody pairs, potentially allowing encoding of multiple, orthogonal signals (Figure 5A). We next tested whether we could report on a native Mtb promoter of interest, the acid-induced promoter Rv1405c, using PPE51-Myc surface display (Rv1405c’::PPE51-Myc). We cultured Rv1405c’::PPE51-Myc Mtb in acidic or neutral minimal medium and stained for Myc displayed by live Mtb along a three-day timecourse. Surface display of Myc was both pH-dependent and time-dependent (Supplementary Figure 7B). However, the kinetics of surface display by fluorescence differed from the induction of fluorescence from Rv1405c’ (Supplementary Figure 8), which may reflect impaired export or increased turnover of surface proteins during acid stress. Nonetheless, it suggests surface display is feasible even under harsher conditions, such as low pH. Surface display could be achieved with other surface protein scaffolds, with PPE36 similarly supporting constitutive and acid-induced display (Supplementary Figure 7C). Taken together, we show that reporting by surface display in Mtb is modular and generalizes to multiple combinations of promoters, scaffolds, and epitope tags.

To determine whether signals from multiple promoters could be simultaneously displayed and detected, we established two dual-reporter systems (dual constitutive hsp60’::PPE51-HA/hsp60’::PPE51-Myc and acid-inducible–constitutive Rv1405c’::PPE51-Myc/hsp60’::PPE51-HA) (Supplementary Figure 9A,D). Compared to single reporters, we observed lower overall expression levels for constitutive dual reporting, suggesting a potential bottleneck associated with displaying multiple scaffolds (Supplementary Figure 9B-C). Acid-induced surface display of Myc and constitutive surface display of HA could be also co-detected after 5 days of culture in acidic minimal medium (Supplementary Figure 9E). Together, this suggests that Mtb surface display can report on multiple aspects of bacterial state using orthogonal promoter-tag pairs.

Our findings suggested that competition in export may contribute to reduced display in multi-display strains. Since PPE51 secretion depends on PE19,^52^ we tested whether surface display was chaperone-limited. We transformed constitutive PPE51-His strains with a low-copy PE19 overexpression vector or an antibiotic-matched empty vector control, then stained OD-matched strains for His. Compared to empty vector, we observed a statistically significant increase in surface display upon chaperone-coexpression (Supplementary Figure 7D). Together, this suggests that surface display levels are chaperone-limited but can be engineered through chaperone co-expression.

### Mtb surface protein truncations generally do not localize to the outer membrane

Given the physiological function of full-length surface proteins, we tested whether truncated Mtb proteins would support outer membrane localization by screening combined hits from tryptic digestion of the surface of intact Mtb,^53^ periplasmically exported fusions from a previous beta-lactamase export screen,^54^ and predicted topologies likely to support C-terminal tagging (Supplementary Figure 10A). Although tryptic digest hits did not entirely overlap with the input beta-lactamase fusion library, we identified multiple shared hits (Supplementary Table 2). However, domain predictions differed between prediction tools, especially for predicted single-pass transmembrane proteins. Incorrect prediction of signal peptides as transmembrane domains is a known issue in topology prediction.^55^

We identified 16 potential protein scaffolds and up to 4 truncations for each protein (Supplementary Table 3), which we C-terminally HA-tagged, expressed, and experimentally validated by HA surface staining. For all truncations tested, we failed to detect HA display, including truncated versions of proteins previously shown to support C-terminal tagging (LprO, Rv1754c, PPE60) and truncations containing up to 75% of the full-length sequence (Supplementary Figure 10B-C). This motivates future studies of outer membrane targeting in Mtb. We propose that mutating away native functions of surface proteins,^52^ combined with co-expression of chaperones, could be a more tractable approach for reducing the impact of surface display on Mtb physiology.

## Discussion

Pathogen-host interactions are associated with heterogeneous pathogen states, host states, and infection outcomes at the population and within-individual level. Understanding these heterogeneous states and how they arise can help us manipulate pathogen-host interactions. However, single-cell methods are limited in their ability to capture pathogen information alongside host information during infection. Here, we repurpose surface display to report on pathogen state and presence axenically and during pathogen-host interactions. We show its extensibility to multiple promoters of interest, display scaffolds, epitope tags, and model and non-model organisms. Toward profiling pathogen-host interactions at finer scales, we demonstrate this method in infected cells down to MOI 1. By engineering bacteria to produce antibody-detectable signals in response to stimuli or transcriptional events, we can detect signals derived from intracellular bacteria by single-cell RNA sequencing without optimizing lysis or bacterial enrichment.

We propose that reporting by surface display is applicable to a range of pathogen-host interaction studies. By using promoters responsive to exogenously provided or endogenously produced small molecules, surface display strains could illuminate host factors underlying heterogeneity in intracellular survival and bacterial metabolic state. Using native promoters, surface display strains could allow joint profiling of bacterial and host transcriptional state at single-cell resolution. Using native secreted proteins and tagging at the genomic locus, surface display strains could report on secretion during infection. By using constitutive promoters and multiple epitope tags, bacterial clinical strains or mutants could be barcoded, as has been previously described for host cells,^27,28^ and used to compare how bacterial variants alter host transcriptional state during infection.

Linear epitope tags are ideal candidates for encoding protein-based signals due to their small size, variety available, and engineerability. Over 20 unique epitope tags exist.^27,28,56^ Novel epitope tags, such as the ALFA tag, can be rationally designed for high-affinity detection.^56^ Combinatorial epitope barcoding strategies, such as adding combinations of antibody-detectable mutations to a single tag^28^ or expressing multiple tags,^27^ can further expand encoding capabilities. Similarly, because antibodies can be conjugated with fluorophores, metals, or DNA, antibody-based detection of bacterial signals should be compatible with a variety of low- and high-dimensional host-focused single-cell assays.

Limits to multi-reporting by surface display are likely organism-dependent. Like fluorescent transcriptional reporters, surface display requires genetic tractability and enables targeted measurement of selected promoters. Where transcriptional modules and representative promoter sequences are defined, such targeted strategies can reduce sequencing cost at the expense of breadth. Engineering surface display systems additionally requires knowledge of outer membrane biogenesis in the organism of interest, and future application to non-model organisms will depend on continued advances in promoter annotation, gene module discovery, and outer membrane proteomics.

Although we focus here on synthetic promoter–epitope fusions, the broader principle is that bacterial state can be encoded in antibody-detectable surface features. As knowledge of state-dependent remodeling of the bacterial surface proteome improves, antibodies against endogenous surface proteins could be used to infer bacterial state without genetic modification. In such contexts, naturally occurring surface markers could substitute for synthetic promoter–epitope constructs, extending this strategy to genetically intractable organisms or clinical isolates.

Reporting by surface display makes signals export-dependent, in addition to translation-dependent. We show this doesn’t create a kinetic delay in all contexts, but does appear to affect total signal for acid-induced Mtb surface display. We hypothesize there is an upper limit to total surface display arising from finite secretion machinery, finite space in the outer membrane, and other mechanisms regulating secretion. We show that export machinery availability can tune secretion, suggesting that signal strength is engineerable. For weak promoters, engineered ribosome binding sites could further boost protein expression.^57^ Statistical methods designed for sparse transcriptomic data sets could also be repurposed to analyze sparse bacterial reporter signals. For experimental conditions that alter protein export or membrane turnover, including a constitutive or small-molecule-inducible surface display tag would allow normalization across experimental and control groups.

Whether fluorescent or surface-displayed, transcriptional reporters strains redirect cellular resources toward transcription and translation of exogenous coding sequences. Surface-displayed transcriptional reporters introduce an extra resource burden on export machinery. Overexpression of native export machinery or introduction of orthogonal export machinery may alleviate this burden, but it is plausible that changes in cellular resources could alter bacterial state or fitness axenically or during pathogen-host interactions. Host state may also be influenced by changes to the bacterial surface proteome. Future work can identify or design minimal scaffolds for resource-efficient, non-immunogenic surface display. Alternatively, future work developing effective antibodies against members of the bacterial surface proteome could be leveraged for dual host-pathogen labeling. This would allow for defining bacterial cell state via surface markers in a similar way to eukaryotic flow cytometry that is also compatible with single-cell transcriptomic methods used here.

We observed heterogeneous HA staining in cells infected with pre-induced, fixed bacteria, despite homogeneous HA staining in the inoculum. By inhibiting phagosomal acidification during infection, we show that HA loss is associated with phagosomal maturation. Non-pathogenic E.coli die within 10 - 60 minutes of infection,^39,40^ but fluorescent proteins expressed in the bacterial cytosol can remain detectable for hours post infection, even when bacteria are pre-killed,^58^ depending upon the specific protein’s pKa and co-factor requirements. As proxies for bacterial transcriptional state, fluorescent transcriptional reporters and epitope tag-based, surface-displayed reporters could provide information with slightly different timescales and biological interpretations. Since loss of HA from mCherry+ iBMMs is bafilomycin-disrupted, it is possible that heterogeneous loss of HA reflects heterogeneous antimicrobial capacity. Coupled measurements of HA and antibody-accessible mCherry could report on bacterial phenotype, i.e. membrane degradation and integrity. However, engineering the display scaffold, linker, and tag to resist degradation or relocalize to a non-degradative compartment could shift surface display toward “recorder”-like behavior and resolve cells containing non-viable, non-intact bacteria.

## Limitations of the study

Surface display–based reporting is inherently targeted, requiring prior selection of promoters and therefore not capturing transcriptome-wide bacterial states. Reporter signal further depends on protein export, surface stability, and host degradative activity, such that loss of signal may reflect bacterial death or degradation rather than transcriptional inactivity, particularly at later timepoints. Display capacity is constrained by secretion and membrane biogenesis machinery, which can limit signal magnitude, multiplexing, and applicability under stress conditions, as observed for Mtb. Extension to non-model or genetically intractable pathogens additionally depends on knowledge of surface protein localization and export pathways, which remain incompletely defined for many organisms. Finally, associations between bacterial reporters and host transcriptional programs are correlative, and establishing causality will require integration with host or pathogen perturbation strategies.

## Acknowledgements

We thank the Bryson lab for helpful discussions. This work was performed in part in the Ragon Institute BSL3 core facility, which is supported by the NIH-funded Harvard University Center for AIDS Research (P30 AI060354). Amy Barczak, Yong Xie, Julie Boucau, Eliane Shwiari, and Stephanie Pringle managed the facility, and Yong Xie assisted with BSL3 flow cytometry. We thank Clifton Barry for sharing PE19 overexpression plasmids and Jeremy Rock for providing advice on strong terminators for Mtb. This work was supported by US NIH grants 1R35GM14290 and R01AI184666 and US Army Grant W911NF-19-D-0001.

## Materials and methods

### Cell and bacterial culture

Immortalized bone marrow macrophages (iBMMs) were grown in DMEM supplemented with 10% FBS. E.coli strains were grown in Miller’s Luria broth. For Mtb, *Mycobacterium tuberculosis* strain H37Rv was grown in 7H9 supplemented with 10% (v/v) OADC, 0.2% (v/v) glycerol, and 0.05% (v/v) tyloxapol. Mtb strains containing plasmids were grown in the presence of appropriate antibiotic: kanamycin (25 ug/mL) or hygromycin B (50 ug/mL).

Minimal medias for H37Rv were prepared using 0.5 g/L asparagine, 1.0 g/L potassium phosphate, 2.5 g/L sodium phosphate, 0.05 g/L ferric ammonium citrate, 0.5 g/L magnesium sulfate, 0.5 mg/L calcium chloride, 0.1 mg/L zinc sulfate, 0.5 ml/L tyloxapol, and 0.2% v/v glycerol, then buffered with 100 mM MES or MOPS for pH 5.7 and pH 7 conditions, respectively.

### Antibodies

Primary antibodies used were: rabbit anti-HA (Cell Signaling Technology, cat#3724); rabbit anti-His (Cell Signaling Technology, cat#12698); rabbit anti-mCherry (Cell Signaling Technology; cat#43590); chicken anti-Myc (exalpha, cat#ACMYC).

Fluorophore-conjugated primary antibodies used were: rabbit anti-HA AlexaFluor 647 (Cell Signaling Technology, cat#37297); rabbit anti-His AlexaFluor 647 (Cell Signaling Technology, cat#14931); rabbit anti-Myc AlexaFluor 647 (Cell Signaling Technology, cat#63730); rabbit anti-FLAG AlexaFluor 647 (Invitrogen, cat#MA1-142-A647).

Secondary antibodies used were: goat anti-rabbit AlexaFluor 488 (Invitrogen, cat#A-11034); goat anti-rabbit AlexaFluor 647 (Invitrogen, cat#A-21245).

### Bacterial staining and flow cytometry

For experiments comparing surface expression across strains, Mtb were inoculated at OD 0.15 two doublings prior to the experiment. For E.coli, overnight cultures were backdiluted to OD 0.1 in LB broth 2 hours prior to the experiment.

For small molecule-inducible expression in E.coli, 200 ng/mL anhydrotetracycline or 4 mg/mL L-arabinose was added to induction conditions for the indicated time. For acid-induced expression in E.coli, strains were grown in LB at pH 4.7 or pH 7 for one hour. For acid-induced expression in Mtb, strains were grown in minimal media buffered at pH 5.7 or at pH 7 for the indicated time.

At endpoint, bacteria were fixed in paraformaldehyde prior to staining, unless otherwise indicated. E.coli strains were fixed for 15 minutes in 2% paraformaldehyde in PBS (v/v), while Mtb strains were fixed 1 hour in 4% PFA:PBS to satisfy inactivation requirements. 100 million bacteria per condition were pelleted, washed, incubated with primary antibody 20 minutes before directly adding the secondary antibody for additional 10 minutes. Primary and secondary antibodies were used at a final concentration of 1:250 in 1% bovine serum albumin in DPBS (w/v). Stained bacteria were washed three times before flow cytometry on a BD FACS Aria cytometer. For experiments with multiple fluorophores, unstained and singly fluorescent strains were used for compensation, except where spectral crosstalk was determined to be minimal.

### Protein isolation and western blotting

For protein isolation from E.coli, bacteria were pelleted, washed once in DPBS, resuspended in 1% SDS in water (v/v). Samples were boiled at 95C for 15 minutes, then clarified by centrifuging at 12,000 xg for 10 mins. Three freeze-thaw cycles were performed on lysates to improve solubilization. Protein concentration in lysates was quantified using a Pierce 660nm assay (ThermoFisher Scientific), then 0.5 ng protein per condition was loaded onto Bis-Tris 4-12% gradient gels, electrophoretically separated, and dry-transferred onto nitrocellulose membranes using the iBlot 2 transfer system. Membranes were blocked for 1 hour at room temperature in TBS blocking buffer (Intercept) and primary stained overnight at 4C at 1:1000 in TBS antibody dilution buffer (Intercept). Primary-stained membranes were washed thrice in TBS-T, secondary stained at 1:10,000 for 1 hour at room temperature, then washed thrice before imaging on a CLx Odyssey imager (LICORbio). Mouse anti-GAPDH (Invitrogen, cat#MA5-15738) was used as a loading control.

### Bacterial transformation

For M.tb transformations, 5 mL mid-log phase bacteria were washed thrice in 10% room temperature glycerol and resuspended in 1/16th of the original volume to generate electrocompetent cells. Electrocompetent bacteria were incubated with 200 ng DNA for 5 minutes at room temperature, then electroporated in a 0.2 cm electrode gap cuvette at 2.5 kV, 25 uF, 1000 ohm. After electroporation, bacteria were recovered in antibiotic-free 7H9 for 1 day before plating on selective 7H10 plates for 17 days.

For E.coli transformations into chemically competent cells, 20 uL NEB DH5a E.coli were transformed by 30 seconds of 42C heat shock, recovered in SOC media 1 hour, then plated overnight on selective LB agar plates. For E.coli transformations into additional strains, mid-log phase E.coli were washed three times with 10% glycerol, halving volume each wash, and stored at -80C until use. Electrocompetent bacteria were incubated with 100 ng DNA for 5 minutes on ice, then electroporated in a 0.2 cm electrode gap cuvette at 2.1 kV, 25 uF, 100 ohm. After recovery in SOC media, bacteria were plated on selective LB plates overnight.

### Infections

E.coli inocula were prepared from washed, mid-log phase bacteria resuspended in pre-warmed DMEM + 10% FBS. iBMMs were infected for 30 minutes, washed three times with DPBS, then incubated 30 minutes in media with or without 200 ng/mL anhydrotetracycline (aTc). Control strains lacking surface expression were also incubated in aTc-containing media to account for background fluorescence.

### Intracellular staining for flow cytometry and single cell RNA sequencing

Cells were detached using 4 mM EDTA in DPBS, then brought to a final concentration of 2% paraformaldehyde (v/v) to fix. After 15 minutes of fixation, fixed cells were pelleted at 1000xg 4C, resuspended in ice cold methanol, and stored at -20C overnight to permeabilize. Permeabilized cells were washed in ice-cold wash buffer (3X saline-sodium citrate, 0.04% ultrapure BSA in water) three times, then blocked for 10 minutes in blocking buffer (3X saline-sodium citrate, 2% ultrapure BSA, 0.1% Tween-20, 0.05% dextran sulfate, 1% Fc Block) before antibody staining with anti-HA, anti-His, or isotype at 2.6 ug/mL.

For single cell RNA sequencing, we added 0.1% recombinant RNAse inhibitor (Takara) and 1mM DTT (Qiagen) to wash and blocking buffers and used a modified fixation protocol. Detached cells were fixed by spiking in fixation buffer (0.33% PFA, 0.01% Tween-20, 0.1% SUPERase IN in PBS) at a ratio of three volumes fixation buffer to one volume cell suspension. After 10 minutes, fixed cells were quenched by addition of 1 M glycine, pelleted, resuspended in ice-cold methanol, and permeabilized overnight at -20℃. For staining, we custom-conjugated the same antibodies with oligos complementary to the 10X Genomics feature barcode capture sequence (Cell Signaling Technology). Stained cells were prepared according to a 10X Genomics standard dual index with feature barcoding protocol, with minor modifications. Stained cells were double-filtered using Flowmi cell strainers (Bel-Art), then pooled and loaded at 6,000 cells / sample in <10 uL wash buffer to limit concentrations of saline-sodium citrate.^59^ During preparation of the antibody library, we used a larger cDNA fraction (10 uL rather than 5 uL) as input for experiments where any lanes were expected to not have epitope tag-derived signal above isotype. Gene expression and antibody libraries were sequenced on a NextSeq 500/550 high output kit with 150 cycles or a NextSeq 2000 XLEAP-SBS P3 kit with 100 cycles. Data was demultiplexed, aligned to mouse genome GRCm38, and quantified using cellranger.

### Estimation of antibody levels from counts

For visualization of background staining, antibody counts were divided by isotype counts, then centered log-ratio normalized. Isotype counts of zero were pseudocounted to 1 to allow division. For normalizing across conditions and defining antibody-positive samples, antibody levels were estimated as counts of antibody-derived tag, then denoised and normalized using dsb,^60^ empty droplets defined by cellranger, and isotype-derived counts. Thresholds for antibody positivity were calculated using ThresholdR^61^ on dsb-normalized antibody levels and k=2 components for bimodal fitting.

### Modeling genes associated with bacterial epitope-derived antibody counts

Gene expression was modeled using a negative binomial generalized linear model in MASS: Gene ∼ dsb(His) + dsb(iso) + log(UMI) + log(genes) + G2M_score + S_score. Only cells in the His-infected condition were considered for this analysis. Genes were modeled as raw counts, and cell cycle scores (G2M, S) were calculated using Seurat’s CellCycleScoring function and mouse orthologs of Seurat cell cycle genes mapped by gprofiler2. For each gene, a null model lacking the dsb(His) feature was also fit, and significance of the dsb(His) feature was calculated by performing an ANOVA on the full and null models and the anova function in the R stats package. ANOVA P-values were adjusted for multiple hypotheses using the Benjamini-Hochberg correction and the p.adjust function in the R stats package.

Multiple cutoffs were used to prioritize significant genes: (1) fold change-based transcriptional variation, (2) percent expression, and (3) p-value of related tests. Transcriptional variation was estimated stringently using Seurat’s FindMarkerGenes function on two cell identities based on thresholded dsb(His) expression and permissively with two cell identities based on K-means=2 clustering of the first 10 principal components of variation. Transcriptional variation was thresholded to require |log2 fold change| > 0.25 between thresholded dsb(His)-defined identities and > 0.5 between K-means–defined identities. Percent expression, as calculated by the FindMarkerGenes function, was thresholded to require >10% expression in at least one identity or a >20% difference in expression level between identities. For stringent identification of His- and phagocytosis-associated genes, we require both adjusted p-values of the two tests performed with FindMarkerGenes < 0.05. For discovery of His- and phagocytosis-associated genes, we require either adjusted p-value < 0.05.

### Data and code availability

Raw data and code are available at: https://github.com/cgunnars/10x_bacterial/. RNA sequencing data generated in this study has been deposited in the Gene Expression Omnibus (GEO) under accession number GSE334578.

---

**Supplementary Figure 1: Surface display is modular with respect to surface protein, and surface signal correlates with intracellular protein expression. A)** aTc-inducible display of His-tagged N-terminal intimin (Neae). Representative of n=3 induction and staining experiments performed on 3 different days. **B)** Schematic of aTc induction timecourse. **C)** Representative surface staining for eCPX-HA at different intervals post aTc addition, representative of n=4 inductions performed on 2 different days. **D)** Representative western blotting for total eCPX-HA at different intervals post aTc addition, representative of n=4 inductions performed on 2 different days. **E)** Correlation between total signal measured by Western blot of cell lysates and surface-displayed signal measured by flow cytometry of intact cells. Total signal is normalized to GAPDH, then maximum epitope tag signal, while surface-displayed epitope tag signal is normalized to maximum epitope tag signal. Error bars = standard deviation, n=3 for pBAD HA/pTet His, n=4 for pTet HA.

**Supplementary Figure 2: Surface displayed epitopes are detectable at a range of MOI conditions. A - B)** iBMMs were infected with fixed surface display strains pre-induced with 200 ng/mL aTc at various MOI for 30 minutes. n=3-4 independent experiments. **A)** Intracellular epitope tag staining and flow cytometry of iBMMs infected with fixed, pre-induced strains at MOI 1 - 50 for 30 minutes prior to staining. **B)** Statistical comparison of HA positivity across MOI 1 - 50, n=5 independent experiments for induced conditions; n=2-3 independent experiments for isotype and uninduced conditions. Multiple comparisons in **B** were calculated based on the arcsin(sqrt(p)) transformation and performed by one-way ANOVA with a post-hoc Holm-Šídák test.

**Supplementary Figure 3 Intraphagosomal retention of surface-displayed signal depends on epitope tag and host degradative capacity. A)** iBMMs were infected with constitutively mCherry-expressing strains 108’::eCPX-HA or 108’::eCPX-His, then stained for HA or His at 30 minute intervals post-phagocytosis. Representative flow plots are shown for 0 and 90 minutes post-phagocytosis, n = 2-4 independent experiments. **B)** Statistical comparison of directly infected (gated on mCherry fluorescence) cells that are also HA- or His-positive at each timepoint. **C)** Representative flow plots of intracellular staining for mCherry at 0 and 90 minutes post-phagocytosis, n = 5 independent experiments. **D)** Statistical comparison of directly infected (gated on mCherry fluorescence) cells that are also mCherry antibody-positive at each timepoint. **E)** Schematic of bafilomycin pretreatment. iBMMs were pre-treated with 75 nM bafilomycin A1 (baf) 1 hour prior to infection. Infections and inductions were performed without continued presence of baf. **F)** Intracellular epitope staining and flow cytometry of baf-pretreated iBMMs infected with live surface display strain and induced post-infection (representative of n = 3 independent experiments). **G)** Statistical comparison of HA positivity, with or without bafilomycin pretreatment.

**Supplementary Figure 4. Timing of aTc addition influences HA positivity. A)** Schematic of aTc withdrawal experiment. **B)** Intracellular epitope tag staining and flow cytometry on iBMM infected with live surface display strains, induced with aTc during phagocytosis or post-phagocytosis. Samples gated using uninfected and isotype-stained samples (representative of n=3 independent experiments) **C)** Statistical comparison of HA positivity in iBMMs with different induction schemes: ++ = induced during phagocytosis and post-infection; +- = induced during phagocytosis, withdrawn post-infection; -+ = induced exclusively post-infection.

Statistical analysis involving percentages were calculated based on the arcsin(sqrt(p)) transformation. Multiple comparisons in **C** were performed by one-way ANOVA with a post-hoc Tukey’s test. Error bars shown represent standard deviations.

**Supplementary Table 1.** Literature evidence for His-associated genes as phagocytosis-associated.

| Gene | Proposed function | Direct evidence regulating phagocytosis? | Associated with phagocytic subsets? |
| --- | --- | --- | --- |
| **Sipa1l1** | Regulates actin assembly during phagocytosis (Xu et al. 2024, JCI Insight) | si-Sipa1l1 increases phagocytosis and actin disassembly (Xu et al. 2024, JCI Insight)<br>Sipa1l1 overexpression decreases phagocytosis and actin polymerization (Xu et al. 2024, JCI Insight) |  |
| **Ckb** | Creatine kinase. Translocates to phagocytic cup, where Ckb enzymatic activity provides ATP to actin assembly (Kuiper et al. 2008, PLoS Biol) | Overexpression of dominant-negative, catalytically dead CKB reduces actin accumulation and phagocytosis (Kuiper et al. 2008, PLoS Biol)<br>Chemical inhibition with cyclocreatine inhibits phagocytosis (Kuiper et al. 2008, PLoS Biol) |  |
| **Cd52** |  | Antibody blockade of CD52 reduces phagocytosis in *ex vivo* monocytes from cirrhosis patients (Geng et al. 2025, J Hepatol) | *Ex vivo* CD52<sup>hi</sup> subsets have higher phagocytic capacity in patients with cirrhosis, compared to CD52<sup>lo</sup> subsets (Geng et al. 2025, J Hepatol)<br>Cd52 transcripts are upregulated in tumor-phagocytic monocyte and macrophage subsets in a cancer mouse model (Gonzalez et al 2023, J Exp Med) |
| **Lamtor2** |  | CRISPR knockout reduces phagocytosis of multiple cargo (Haney et al 2018, Nat Genet; Essletzbichler et al 2023, Life Sci Alliance) | Lamtor2 transcripts are upregulated in tumor-phagocytic macrophage subsets in a cancer mouse model (Gonzalez et al 2023, J Exp Med) |
| **Ifitm3** |  |  | Ifitm3 transcripts are upregulated in tumor-phagocytic monocyte subsets in a cancer mouse model (Gonzalez et al 2023, J Exp Med) |
| **Anxa1** | Annexin A1. Translocates to the phagocytic cup, controls F-actin association with the phagosome (Patel et al 2011, J Cell Sci) | Macrophages isolated from whole-mouse Anxa1<sup>-/-</sup> knockout have impaired phagocytosis of multiple cargo (Yona et al 2006, Br J Pharmacol)<br>si-Anxa1 RAW.647 macrophages have impaired phagocytic cup formation and F-actin-phagosome association (Patel et al 2011, J Cell Sci)<br>Antibody blockade of Annexin A1 impairs F-actin-phagosome association (Patel et al 2011, J Cell Sci) |  |

**Supplementary Figure 5 Transcriptional heterogeneity is present at early timepoints and associated with uptake of HA-positive bacteria. A - C)** Transcriptional clusters in iBMMs infected with (**A**) pre-induced Tet::eCPX-HA E.coli (**B**) uninduced Tet::eCPX-HA, and (**C**) uninfected iBMMs at 0 minutes post-phagocytosis. Dsb-normalized HA counts are plotted for each cluster. **D)** Expression of stringently His-associated genes in samples infected with pre-induced or uninduced bacteria and in uninfected samples. **E)** Comparison of His-associated marker expression in 108::eCPX-His–infected samples at 90 minutes post-phagocytosis and uninduced, infected or uninfected samples at 0 minutes post-phagocytosis. Fold change between transcriptional clusters is plotted for genes significantly associated with His levels using discovery cutoffs. Labels are shown for genes that meet stringent cutoffs. **F)** Comparison of cluster markers between pre-induced, uninduced, and uninfected samples. Fold change between transcriptional clusters is plotted for significant cluster markers in the pre-induced condition.

Comparisons in **A-C** are calculated using a two-sided Kolmogorov-Smirnov test, and goodness of fit and correlations in **E-F** are calculated using linear regression and Pearson’s correlation test.

**Supplementary Figure 6 Mouse macrophages heterogeneously express phagocytosis-associated genes at the single-cell level. A - B)** Expression of stringently His-associated genes is plotted for unstimulated, wildtype BMDMs collected by (**A**) Jani et al and (**B**) Muñoz-Rojas et al. Genes not meeting an expression/capture threshold of >20% cells with non-zero counts within a given condition are omitted.

**Supplementary Figure 7 Reporting by surface display in Mtb is engineerable and generalizes to different promoters, surface proteins, and tags. A)** Surface staining of epitope-tagged PPE51 expressed under the constitutive hsp60 promoter. Representative of n=3 independent staining experiments performed on 3 different days for His, Myc, and HA; n=2 for V5. **B)** Surface staining of Rv1405c’:PPE51 during culture in low pH or neutral minimal medium over 5 days. Representative of n=3 inductions performed on 2 different days. **C)**. Surface staining of constitutive PPE36 display strains and acid-inducible PPE36 display strains after 5 days of induction. Representative of n=2 independent staining experiments performed on two different days for constitutive display and n=2 independent staining experiments performed on the same day for acid-inducible display. **D)** Surface display of constitutive PPE51-His in strains co-transformed with a low-copy vector constitutively expressing PE19 or empty vector, representative of n=3 independent inductions performed on 3 different days. His gMFI is quantified.

Statistical analysis involving multiple comparisons **(B)** and **(F)** was performed by one-way ANOVA with a post-hoc Tukey’s test, while single comparisons **(J)** were performed by ratio paired t-test.

**Supplementary Figure 8 Fluorescent reporters of Rv1405c are induced within a day of acidic culture.** Rv1405c’::mNeonGreen reporter strain was induced in pH 5.7 or pH 7 minimal media with glycerol as a carbon source, and fluorescent signal was measured by flow cytometry on live bacteria.

**Supplementary Figure 9 Surface display in Mtb generalizes to dual promoter-tag display, but is less efficient. A)** Schematic of dual surface display vector. **B)** Dual surface display of PPE51-Myc and PPE51-HA using two constitutive hsp60 promoters. Representative of n=3 independent staining experiments performed on 2 different days. **C)** Co-staining of HA and Myc in dual display constructs, single PPE51-HA or PPE51-Myc constructs, and wild-type Mtb. **D)** Schematic of dual, acid-inducible and constitutive surface display. **E)** Representative co-staining for acid-inducible Myc and constitutive HA in dual display strains, exposed to acidic or neutral minimal medium for 5 days, n=2 independent staining experiments performed on the same day

**Supplementary Figure 10 Truncations of the surface protein lprO ablate surface localization. A)** Strategy for identifying protein truncations that support outer membrane localization. Probabilities and domain predictions for full-length lprO using TMHMM 2.0. **B)** Outer membrane localization of full-length Rv1745c and PPE60, representative of n=4 independent staining experiments performed on 2 different days. **C)** Surface staining and flow cytometry of lprO truncations, relative to full-length lprO, wild-type H37Rv as a negative staining control, and constitutively expressed PPE51 as a positive staining control, representative of n=2 independent staining experiments performed on 2 different days.

**Supplementary Table 2.**
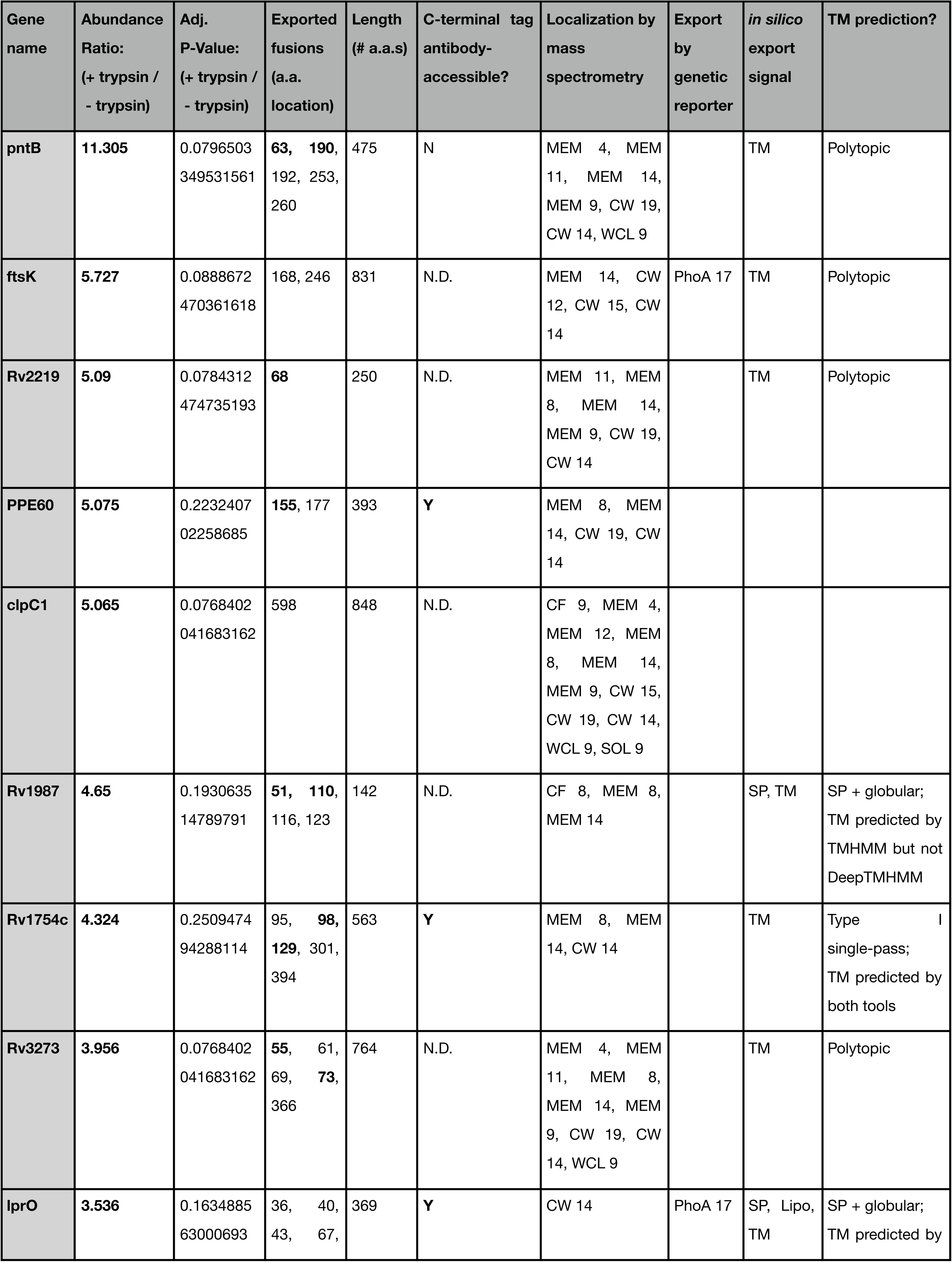
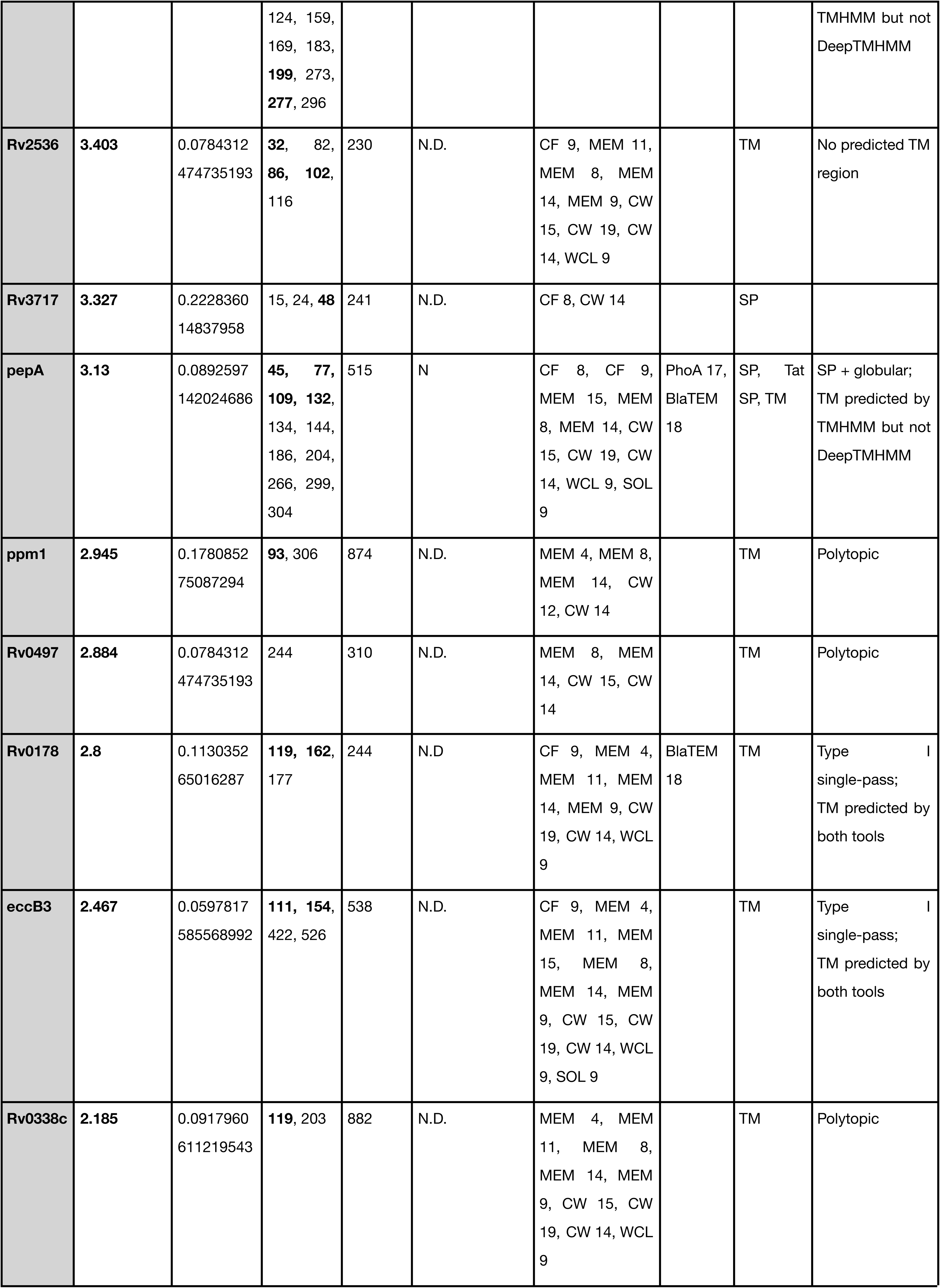
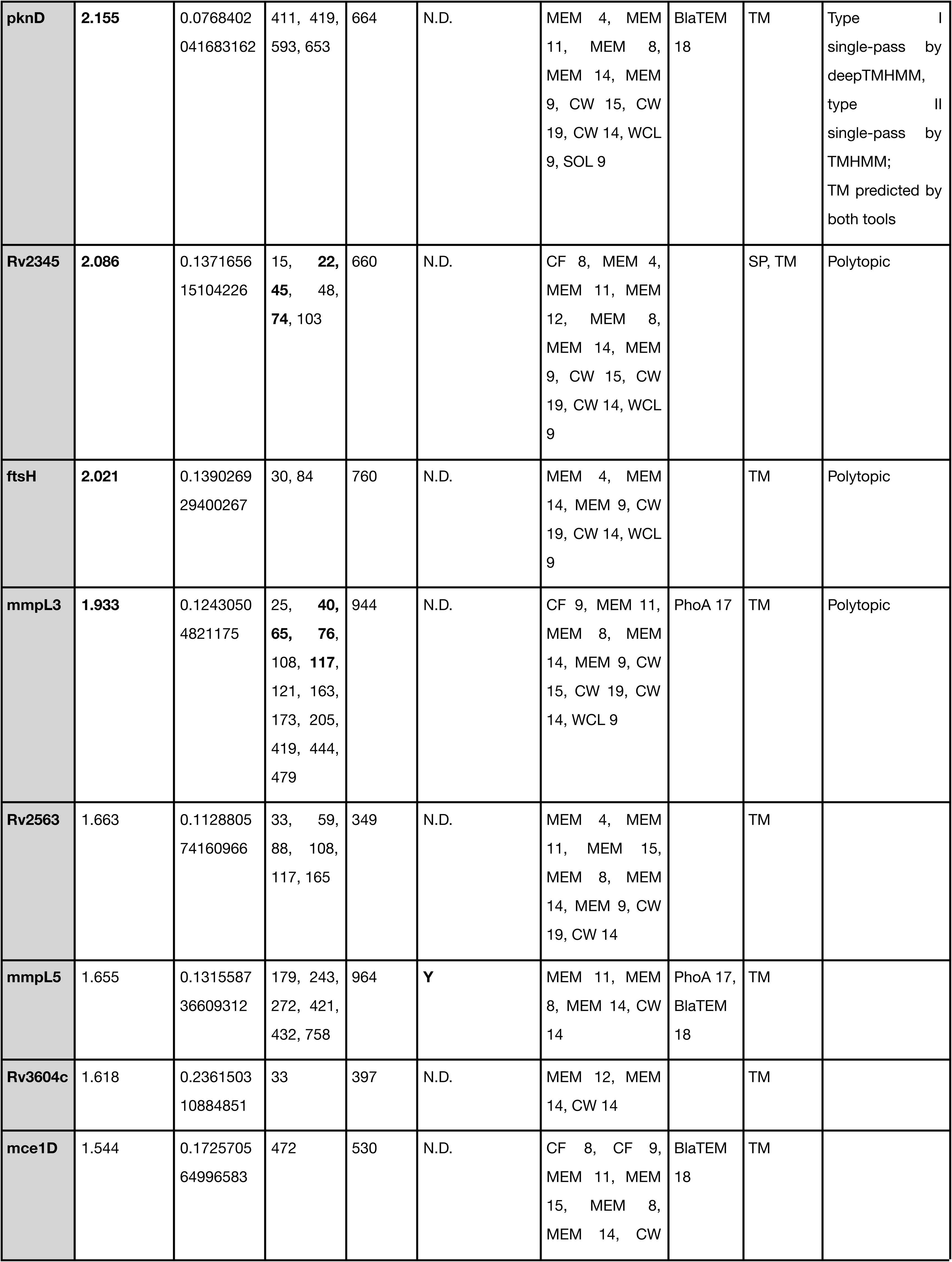
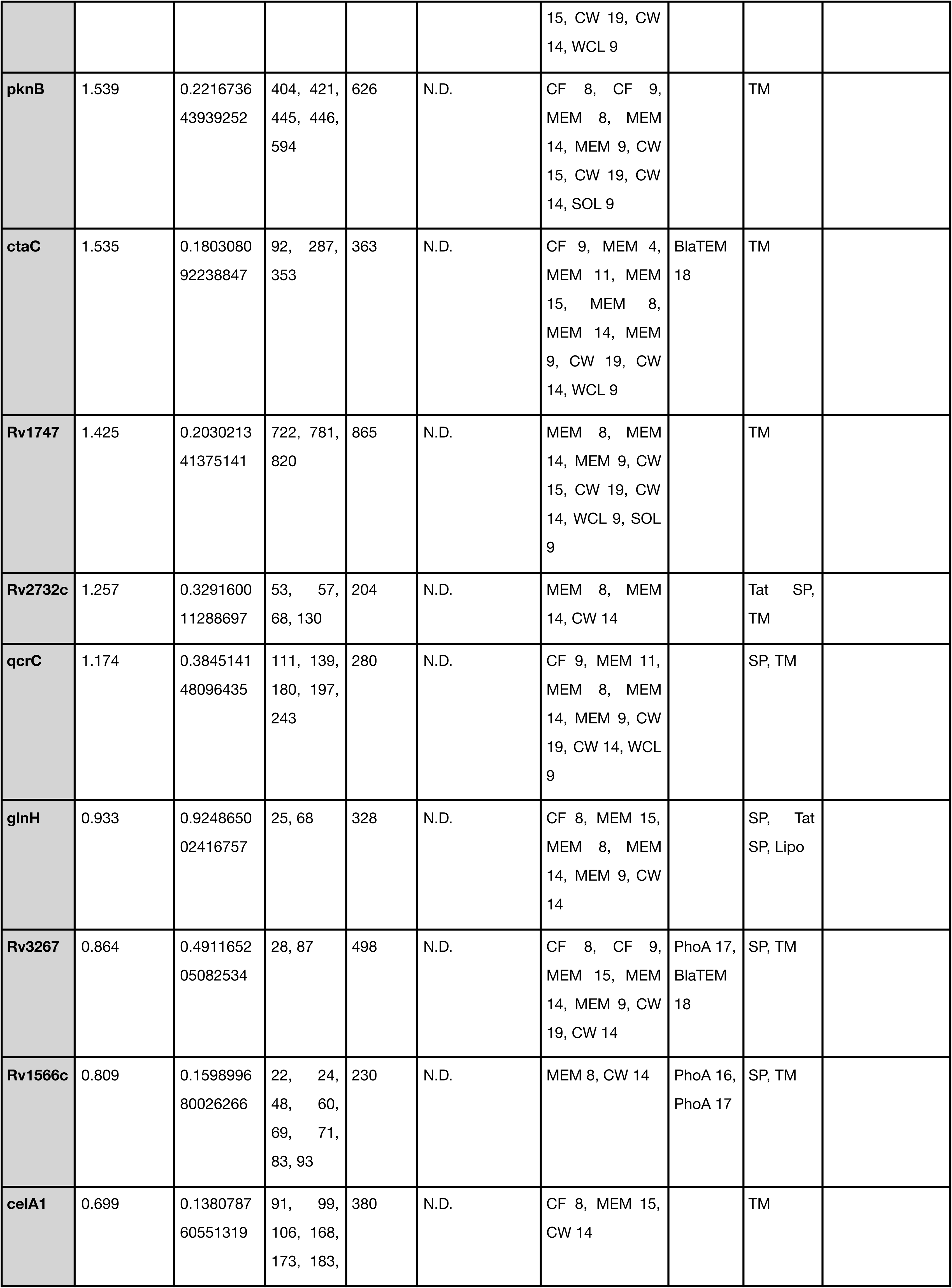
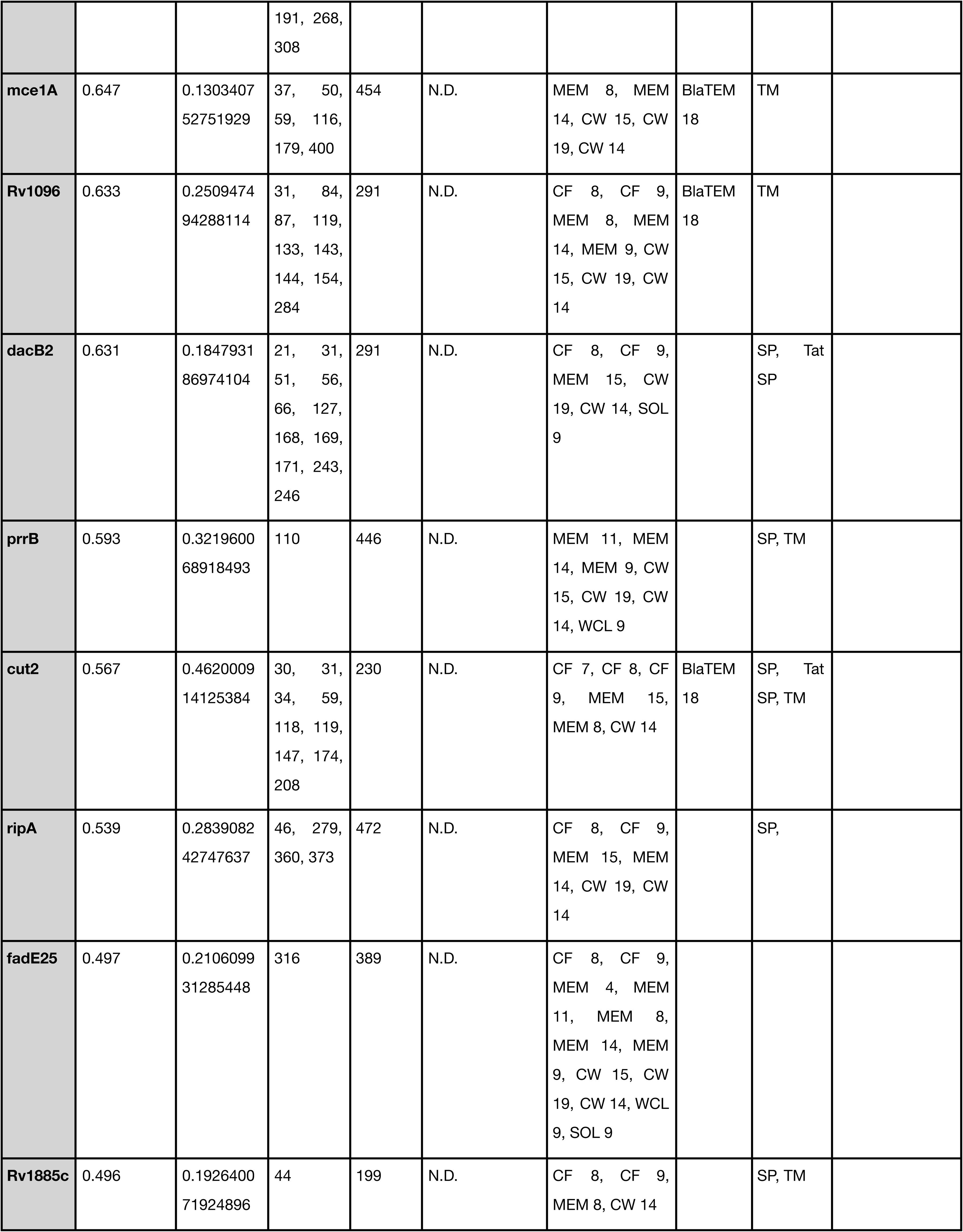
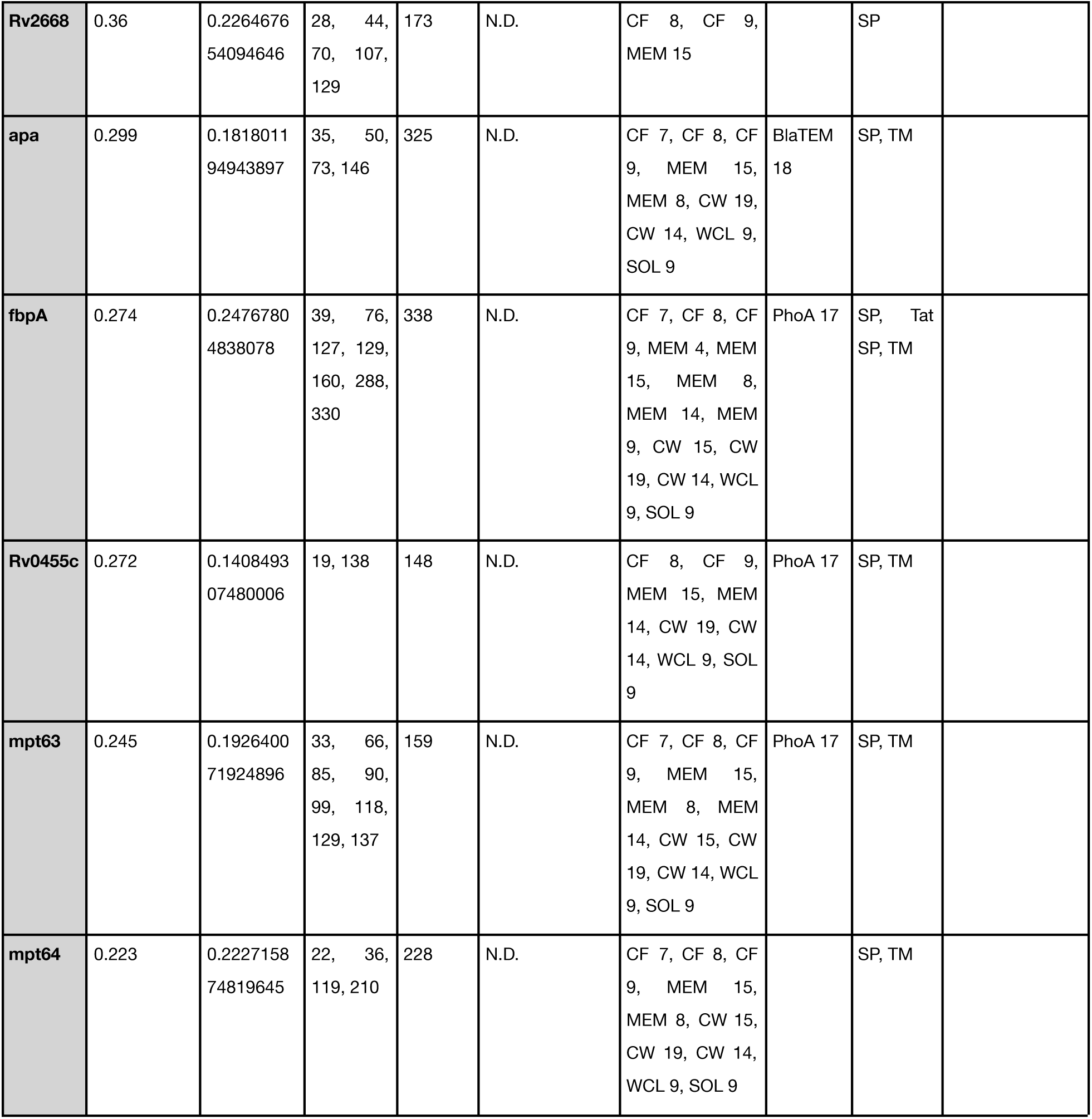
Compiled trypsin accessibility, beta-lactamase export, and transmembrane domain prediction data for Mtb proteins. Proteins for which trypsin accessibility data and beta-lactamase export data was available are tabulated. Columns 1 - 2 are reproduced from Lepe et al 2025, columns 3 - 7 are reproduced from Perkowski et al 2017. Column 8 contains updated predictions of transmembrane topology calculated using TMHMM 2.0 or deepTMHMM 1.0 and was generated as original work. For each protein, the abundance and p-value of trypsin-accessible fragments relative to a no-trypsin control is reported. N-terminal fragments of each protein that support beta-lactamase secretion are listed, and fusions tested for each protein are **bolded**.

**Supplementary Table 3.** All tested truncations. HA-tagged fusions of the first n amino acids of each gene were cloned, transformed in H37Rv Mtb, and stained for surface HA in n=2 independent experiments.

| Gene name | Tested truncations (first N a.a.s) | Untruncated length (# a.a.s) |
| --- | --- | --- |
| pntB | 63,190 | 475 |
| Rv2219 | 68 | 250 |
| PPE60 | 155 | 393 |
| Rv1987 | 51, 110 | 142 |
| Rv1754c | 98, 129 | 563 |
| Rv3273 | 55, 73 | 764 |
| lprO | 199, 277 | 369 |
| Rv2536 | 32, 86, 102 | 230 |
| Rv3717 | 48 | 241 |
| pepA | 45, 77, 109, 132 | 515 |
| ppm1 | 93 | 874 |
| Rv0178 | 119, 162 | 244 |
| eccB3 | 111,154 | 538 |
| Rv0338c | 119 | 882 |
| Rv2345 | 22, 45, 74 | 660 |
| mmpL3 | 40, 65, 76, 117 | 944 |

## Notes

### Competing Interest Statement

The authors have declared no competing interest.

